# The early embryonic transcriptome of a Hawaiian *Drosophila* picture-wing fly shows evidence of altered gene expression and novel gene evolution

**DOI:** 10.1101/2021.10.29.466520

**Authors:** Madeline Chenevert, Bronwyn Miller, Ahmad Karkoutli, Anna Rusnak, Susan Lott, Joel Atallah

**Author notes:** **Madeline Chenevert:** Hayward Genetics Center, Tulane University School of Medicine, 1430 Tulane Avenue, SL-31, New Orleans, LA 70112. **Anna Rusnak:** Center for Biomedical Engineering, Brown University, Box A-2, Arnold Lab, 91 Waterman Street, Suite 211, Providence, RI 02912. **Ahmad Karkoutli:** LSUHSC School of Medicine, 1901 Perdido St, New Orleans, LA 70112.

## Abstract

A massive adaptive radiation on the Hawaiian archipelago has produced approximately one quarter of the fly species in the family Drosophilidae. The Hawaiian *Drosophila* clade has long been recognized as a model system for the study of both the ecology of island endemics and the evolution of developmental mechanisms, but relatively few genomic and transcriptomic datasets are available for this group. We present here a differential expression analysis of the transcriptional profiles of two highly conserved embryonic stages in the Hawaiian picture-wing fly *Drosophila grimshawi*. When we compared our results to previously published datasets across the family Drosophilidae, we identified cases of both gains and losses of gene representation in *D. grimshawi*, including an apparent delay in Hox gene activation. We also found high expression of unannotated genes. Most transcripts of unannotated genes with open reading frames do not have homologs in non-Hawaiian *Drosophila* species, although the vast majority have sequence matches in other genomes of the Hawaiian picture-wing flies. Some of these genes may have arisen from non-coding sequence in the ancestor of Hawaiian flies or during the evolution of the clade. Our results suggests that both the modified use of ancestral genes and the evolution of new ones may occur in rapid radiations.

**RESEARCH HIGHLIGHTS:** The early embryonic transcriptome of the Hawaiian fly *Drosophila grimshawi* shows a loss of expression of conserved Stage 5 genes, including the Hox genes

The de novo evolution of embryonically expressed genes may be occurring in the Hawaiian *Drosophila* lineage

**AUTHORS’ STATEMENT:** This paper is not being considered for publication elsewhere. This study formed part of Madeline Chenevert’s M.S. thesis.

## INTRODUCTION

It is estimated that 1,000 of the world’s 4,000 Drosophilid fly species are endemic to the Hawaiian archipelago (O’Grady & DeSalle, 2018). Comprising the largest, and arguably the most diverse, radiation within the family, Hawaiian *Drosophila* flies (Figure 1) show extensive phenotypic variation, with morphological modifications at every stage of the life cycle (Craddock et al., 2018; Sarikaya et al., 2019). Flies of the iconic Hawaiian picture-wing clade (Magnacca & Price, 2015), for example, show highly varied wing patterns that may be sexually monomorphic or dimorphic (Edwards et al., 2007). Other clades are named after characteristic modifications to the tarsus (Lapoint et al., 2009, 2014) or mouth parts (Magnacca & Grady, 2009), both specific to male flies. In addition to changes to the adult body plan, the eggs display a vast range of variation in size, structure, and filament number (Kambysellis et al., 1980; Kambysellis & Heed, 1971). This phenotypic diversity raises the question of whether there is similar variation in gene expression, and whether the evolution of novel genes (Van Oss & Carvunis, 2019), or the re-use of ancestral ones (Carroll, 2008), underlies the diversification of the clade.

**Figure 1.**
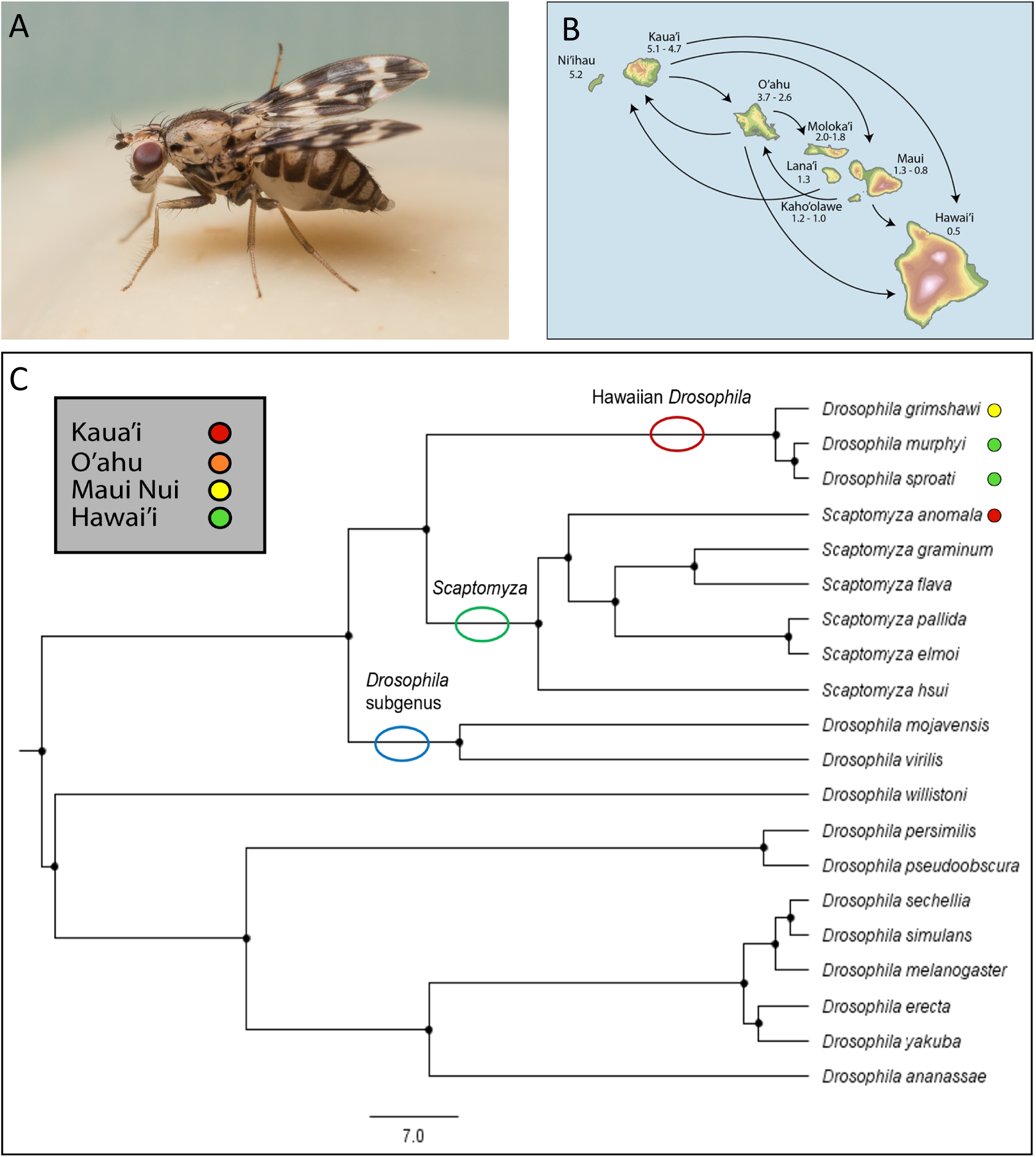
Hawaiian *Drosophila*. A) *Drosophila grimshawi*. B) A map of the Hawaiian islands, with arrows showing inferred colonization events. Kaua’i is the oldest of the larger islands and Hawai’i the youngest. Mau’i Nu’i consists of Moloka’i, Lana’i, Mau’i and Kaho’olawe. Most colonization events occurred from older (northern) to younger (southern) islands, although there were also cases of reverse colonization. C) A phylogeny of 20 Drosophilidae species. The Hawaiian *Drosophila* lineage is a sister-group to *Scaptomyza*. The ancestor of both clades was probably Hawaiian, but *Scaptomyza* dispersed globally, while the Hawaiian *Drosophila* remained confined to the archipelago. The phylogeny is based on multiple studies (Katoh et al., 2017; Magnacca & Price, 2015; Ometto et al., 2013) and divergence times gleaned from TimeTree (Hedges et al., 2015; Kumar et al., 2017) were used in its construction.

Transcriptional profiling of Hawaiian flies, however, has been limited. While recent studies have carried out functional genomics in adult stages of picture-wing flies (Kang et al., 2016), gene expression levels in the embryo have not previously been analyzed. Hawaiian flies are notoriously difficult to culture (Montgomery, 1975), and the life cycle is extended relative to common laboratory species such as *Drosophila melanogaster*, complicating both the collection of samples and the interpretation of results from timed specimens.

Early embryogenesis, which constitutes the opening act in an organism’s life (Tadros & Lipshitz, 2009), is the basis for all subsequent development, and perhaps the most critical period to first consider when analyzing a species with a dearth of transcriptomic data that is difficult to culture. Across Drosophilidae, early embryos pass through comparable developmental stages (Kuntz & Eisen, 2014). We have previously shown (Atallah & Lott, 2018) that gene transcript levels are highly concordant across species at two early embryonic stages, stage 2 (St2) and late stage 5 (St5) (Figure 2). The first of these stages precedes the maternal-zygotic transition, while at stage 5 zygotic expression is underway and many maternal transcripts have been degraded (Figure 2). While most genes are represented at both stages, those that are St5-only (zygotically transcribed with no maternal contribution) are strongly conserved across large phylogenetic distances (Atallah & Lott, 2018).

**Figure 2.**
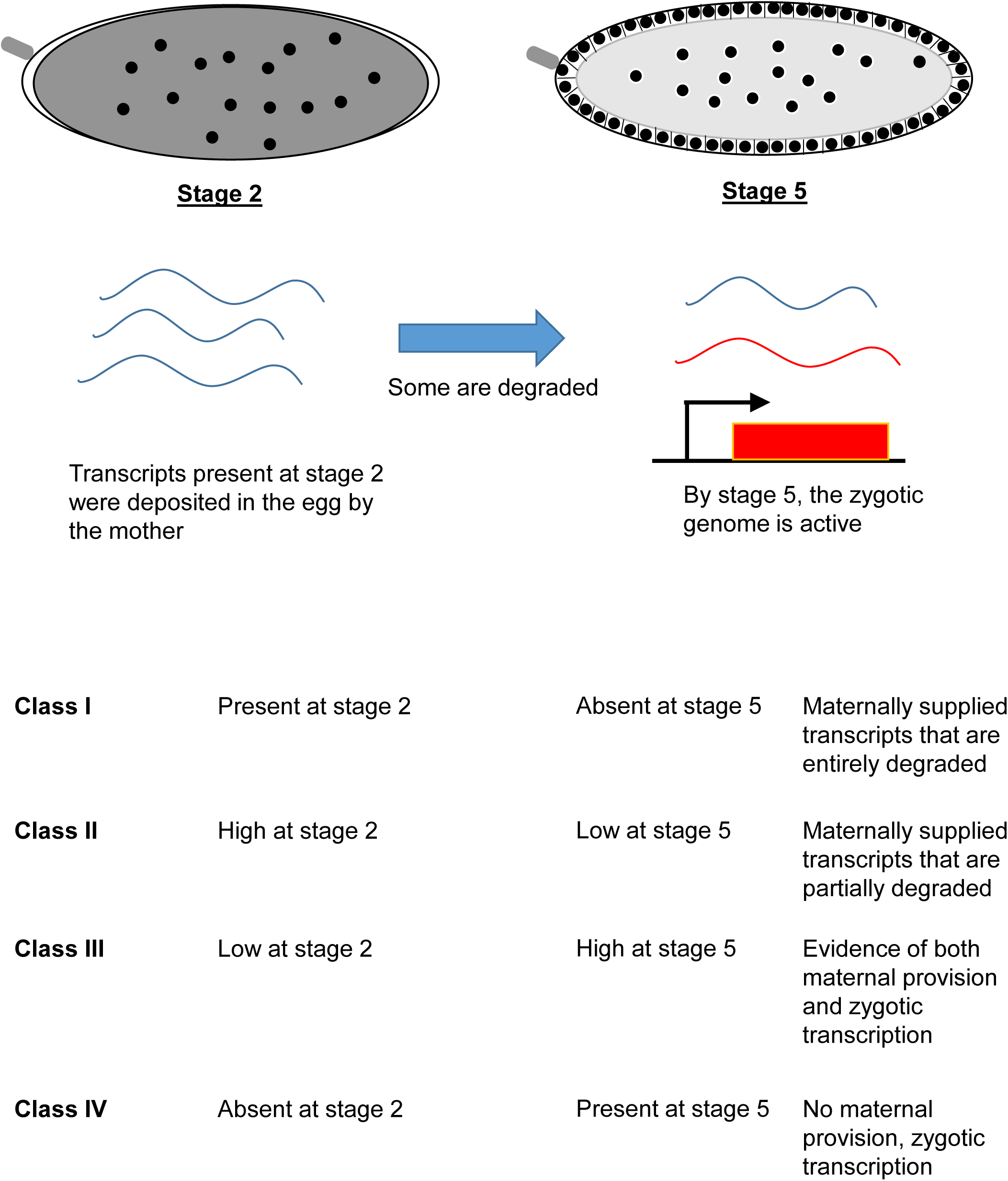
The maternal-to-zygotic transition (MZT). The zygotic genome is silent in the early embryo, which relies on transcripts that were deposited by the mother and translated after egg activation to jump-start development. Many of these transcripts are degraded. By stage 5, the embryonic transcriptome consists of both maternally deposited mRNAs that have not been degraded and zygotically transcribed mRNAs.

Here, we present results of a single-embryo RNA-Seq analysis of these critical stages in the Hawaiian picture-wing fly *D. grimshawi*. We compare the findings to our previously published data on other *Drosophila* flies. We find that *D. grimshawi* shows extensive loss of gene transcript representation at zygotic genome activation (stage 5) relative to non-Hawaiian species. Notably, in sharp contrast to outgroup clades, we find no evidence that the Hox genes, the downstream component of the antero-posterior segmentation cascade, are activated by stage 5. We further find instances of early embryonic mRNA representation of gene orthologues in this picture-wing fly which are not seen in other species until later stages in development. Finally, we conduct an analysis of unannotated genes in *D. grimshawi* that are represented in the early embryo, and provide evidence that many of them are taxonomically restricted. Some of these genes appear to have been generated de novo from non-coding sequence, either in the Hawaiian *Drosophila* proper or in the ancestor of Hawaiian *Drosophila* and its sister-clade, *Scaptomyza* (Figure 1), which is believed to have originated in Hawaii before diversifying globally (Lapoint et al., 2013; O’Grady & DeSalle, 2008).

## MATERIALS AND METHODS

### Drosophila grimshawi husbandry

Upon receiving the *D. grimshawi* stock (Stock Number 15287-2541.00) from the San Diego *Drosophila* Species Stock center (currently the National *Drosophila* Species Stock Center at Cornell University, https://www.drosophilaspecies.com/), the flies were maintained in accordance with a modified version of stock center protocols. The line we used was a replacement donated by Dr. Ken Kaneshiro from the same stock that was used in sequencing the *D. grimshawi* genome for the 12 *Drosophila* Genomes Project (Clark et al., 2007). The flies were kept in vials of Wheeler-Clayton food (Wheeler & Clayton, 1965). To counter the high humidity of New Orleans, which could have caused the flies to stick to the food, Wheeler-Clayton food was prepared using the maximum recommended amount of agar (14.5 g per liter of water). The cooked potato media was prepared with 456 g of filtered water for every 100 g of powdered mix to ensure a consistency that prevented food from falling during transfers. High doses of ethanol and propionic acid (6.5 ml per liter of water) were also used in the top layer recipe to prevent mold growth.

Vials were papered with Kimwipes, and flies were transferred to new vials twice per week. We found that maintaining stocks at a density of 12-16 flies per vial maximized fecundity while reducing death due to overcrowding. Vials containing third instar larvae were transferred to jars to pupate. The jars contained equal volumes of oolite and aragonite sand, approximately 2 cm deep, and were covered with two layers of heavy-duty paper towels. The sand was moistened with water from a spray bottle once or twice per week, with the aim of keeping the sand and towels damp while avoiding puddles. Excess water was absorbed with cellulose acetate plugs (“flugs”, Genesee Scientific), which were then disposed of.

After the first eclosions, the vials with larvae were moved to a new sand jar, and a small petri dish containing the top layer of the Wheeler-Clayton food was added to the jar to feed new flies as they emerged. Initially, we rapidly transferred the recently-eclosed flies to vials to expand our population, but we observed that they often became stuck in the paper or condensation due to the high ambient humidity in New Orleans. To avoid losing adults before they reached sexual maturity, the flies were maintained in the jar for two weeks, then moved to large egg collection cages. The cages were placed atop petri dishes containing a base of apple-agar media, combined with 2 mL of melted Wheeler-Clayton food, providing nutrient-rich media to encourage egg-laying. These Wheeler-Clayton pellets, containing the eggs, were transferred from the apple media with a spatula, washed with water or a 50% solution of apple cider vinegar to inhibit mold, cut into multiple pieces with a spatula and mixed into vials of Wheeler-Clayton food. Most vials prepared in this manner produced dozens of larvae and little to no mold.

### Egg collection

Single embryo collection and staging were carried out as described previously (Atallah & Lott, 2018), with a few modifications. Briefly, flies were maintained at room temperature in an egg collection cage lined with a petri dish filled with apple-agar media, with 2 ml of a paste of live yeast in the center of the plate. Since stage 2 embryos can confused with later stages, the cages were timed to ensure all eggs were laid within one hour. The flies were maintained on live yeast for an hour, after which the plate was discarded and replaced with a new one. Stage 5 embryos were collected from these plates, and Stage 2 embryos were only collected after plates were replaced once more.

Eggs were dechorionated with a 50% bleach solution. Unlike smaller eggs of non-picture wing species, which could be dechorionated after less than two minutes in bleach, dechorionation of *D. grimshawi* eggs could take upwards of three minutes. The process was usually halted when the filaments were completely dissolved and no longer visible. Eggs were rinsed with deionized water for at least 30 seconds. The dechorionated eggs were then transferred to a drop of halocarbon oil (Sigma) on a microscope slide. The slide was placed on a dissection scope over refracted light. Individual embryos were selected using a set of morphological characteristics (Bownes, 1975). St2 embryos were identified by the empty poles on each end of the egg and their lack of visible cell membranes and evagination. St5 embryos were collected at the end of stage 5, after cellularization and before gastrulation. Traits required for St5 embryos were 2 layers of well-defined cell membranes and formation of the pole cells. Embryos showing damage or any invagination were rejected.

### Embryo processing and RNA extraction

A 1.5 mL microcentrifuge tube was filled with 800 μl of Trizol reagent (Ambion) and labelled with the sample number. Each embryo was placed on the corner of a clean microscope cover slip. Using a fine paintbrush, the embryo was transferred to a new location on the slide to remove excess halocarbon oil. After repeating this process two or three times, the embryo was positioned in the center of the slide for lysing. 3 μl of Trizol were pipetted from the labelled 800-ul aliquot and placed in a droplet on top of the embryo. The embryo was then lysed with a sterile, 30-gauge medical lancet (Reli-On Ultra-Thin). Embryos were left to dissolve in the Trizol for 5 minutes. After the embryo had completely dissolved, an additional 3 ul of Trizol from the tube were added to the slide, and all 6 ul were pipetted and transferred to the tube containing the remaining 794 μl, which was mixed by pipetting multiple times. 6 microliters were again removed from the same tube and pipetted onto the place of the embryo, re-collected and returned to the aliquot. This rinsing step was repeated twice to ensure thorough collection of all embryonic tissues. Labelled samples were stored at −80°C until RNA isolation.

RNA was extracted per the manufacturer’s instructions using a Trizol phenol-chloroform extraction (Invitrogen) in which the method was modified to accommodate an initial volume of 800 ul of Trizol. During RNA precipitation, 10 μL of 20 μg/μl glycogen (Invitrogen™ UltraPure™ Glycogen) was used, and the samples were spun in an Eppendorf 5424 R centrifuge at 21130 RCF (maximum speed) for a full 60 minutes. If a sample showed no visible pellet at this point, it was spun for another 30 minutes. After resuspension in UltraPure water, 15.5 μl were kept for processing, and 3 aliquots of 1.5 μl each were taken for quality analysis.

### RNA Quality analysis

RNA integrity was assessed on an Experion bioanalyzer using a Bio-Rad Experion RNA High-Sensitivity kit or an Agilent 2100 bioanalyzer using a Bioanalyzer RNA 6000 Pico assay (Agilent). RNA concentrations were measured using a Qubit 4 fluorometer with a Qubit high-sensitivity RNA assay.

### cDNA libraries

To remove any DNA contamination before library construction, a Turbo DNA-Free kit was used according to the manufacturer’s directions before immediately continuing to library construction using the NEBNext Ultra RNA Library Prep kit for Illumina. 15 PCR cycles were used in the final enrichment step of library generation. cDNA quality was assessed using an Agilent 2100 Bioanalyzer with a High-Sensitivity DNA Kit. The samples showed main peaks around 300 base pairs, as expected.

### Sequencing

The four highest quality libraries from each stage were sent to Novogene for sequencing. Only libraries from samples showing minimal RNA degradation were selected. In the final analysis, 3 libraries from each stage were used. One library from stage 2 was not used due to low overall read mapping rate (less than 65%) while another library from stage 5 appeared to have been mislabeled. The files containing the reads are publicly available at NCBI’s Sequence Read Archive (BioProject Accession Number PRJNA771180).

### Transcriptome mapping, assembly and differential expression

Bioinformatic analysis was carried out as described previously (Atallah & Lott, 2018). Briefly, adapters were removed using Cutadapt (Martin, 2011) and reads were trimmed and filtered for quality. We used the Tuxedo suite (Trapnell et al., 2012) for transcriptomic analysis. Although much faster alignment-free methods have become more popular in recent years, we chose the Tuxedo suite both to allow the identification of novel isoforms and genes and for straightforward comparison with our previous analyses of equivalent stages in non-Hawaiian species. We aligned the reads, using Tophat2 (D. Kim et al., 2013), to the GCF_000005155.2_dgri_caf1 *D. grimshawi* NCBI reference genome. Cufflinks (Trapnell et al., 2013) was used with the -N upper-quartile normalization option. Gene expression levels were determined by combining the expression of all gene isoforms. Differential expression was determined using CuffDiff. *D. melanogaster o*rthologs were assigned using the Flybase (Larkin et al., 2021) orthology table. An FPKM threshold of 1 was used, as employed previously (Atallah & Lott, 2018).

### Gene Ontology (GO) enrichment analysis

GO enrichment of genes with identified one-to-one *D. melanogaster* orthologs was assessed using DAVID (Huang et al., 2009a, 2009b). In cases where Cufflinks combined two annotated genes, both genes were considered differentially expressed in the GO analysis. The background list in all cases was the set of all genes represented in the *D. grimshawi* embryo, unless otherwise specified. The R package GOPlot (Walter et al., 2015) was used to generate graphical representations of gene enrichment.

### Unannotated gene analysis

Transcripts of genes that were identified by Cufflinks but had no annotation in the NCBI genome were searched for open reading frames using TransDecoder (Haas et al., 2013). Those with complete open reading frames of at least 50 amino acids were then compared to both the sequenced genomes of 11 other *Drosophila* species and previously generated embryonic transriptome assemblies of the same species (Atallah & Lott, 2018; Combs & Eisen, 2013; Lott et al., 2014; Paris et al., 2015). For cases where no ortholog was identified using BLASTP (Altschul et al., 1990), we used TBLASTN to search for matches in the other *Drosophila* genomes, then analyzed these matches for ORFs using TransDecoder. Alignments were generated using CLUSTALW (Thompson et al., 2003).

## RESULTS

### Gene representation in the early *D. grimshawi* embryo

We found a total of 18,617 transcripts to be represented in the early embryo above our established threshold (see Materials and Methods). These transcripts belonged to 8,233 genes. Genes with one-to-one orthologs in *D. melanogaster* or other species are shown in Table S1. Using insights obtained from previous analyses, we identified four classes of genes (Figure 2), discussed below. We used the R package GOPlot (Walter et al., 2015) to graphically represent the results of our GO analysis as concentric circular plots (Figures 3 and 4).

**Figure 3.**
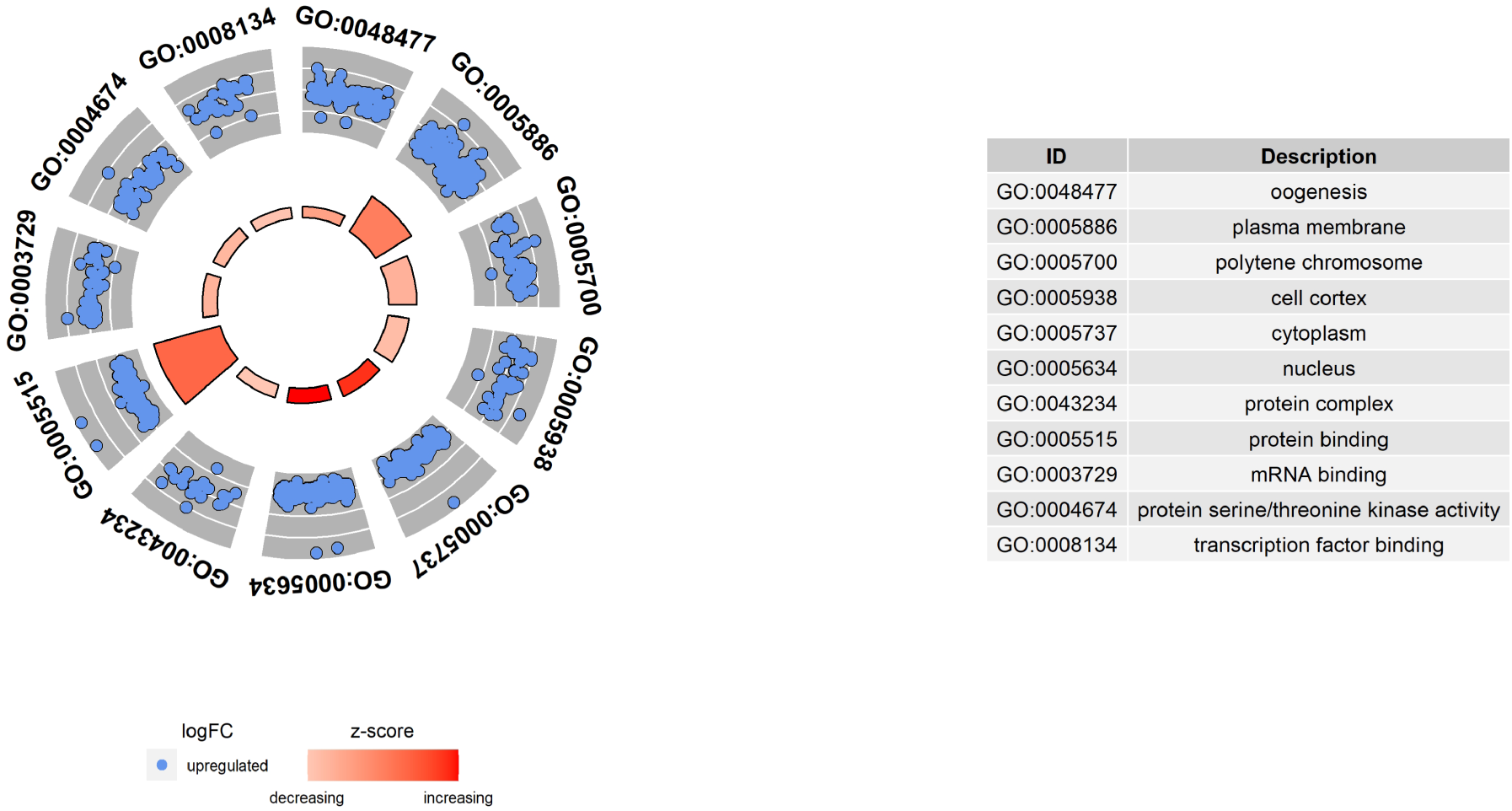
Gene Ontology (GO) analysis of genes that are represented at both stages but are higher at St5 (Class 3). The height of the bars in the inner circle represents the Benjamini-corrected enrichment P-value of the associated term, and the color represents the “z-score”, in this case the square root of the number of genes mapping to the term. The outer circle shows the log of the fold enrichment (St5/St2) of genes mapping to each term. GOplot (Walter et al., 2015) was used to generate the plot.

**Figure 4.**
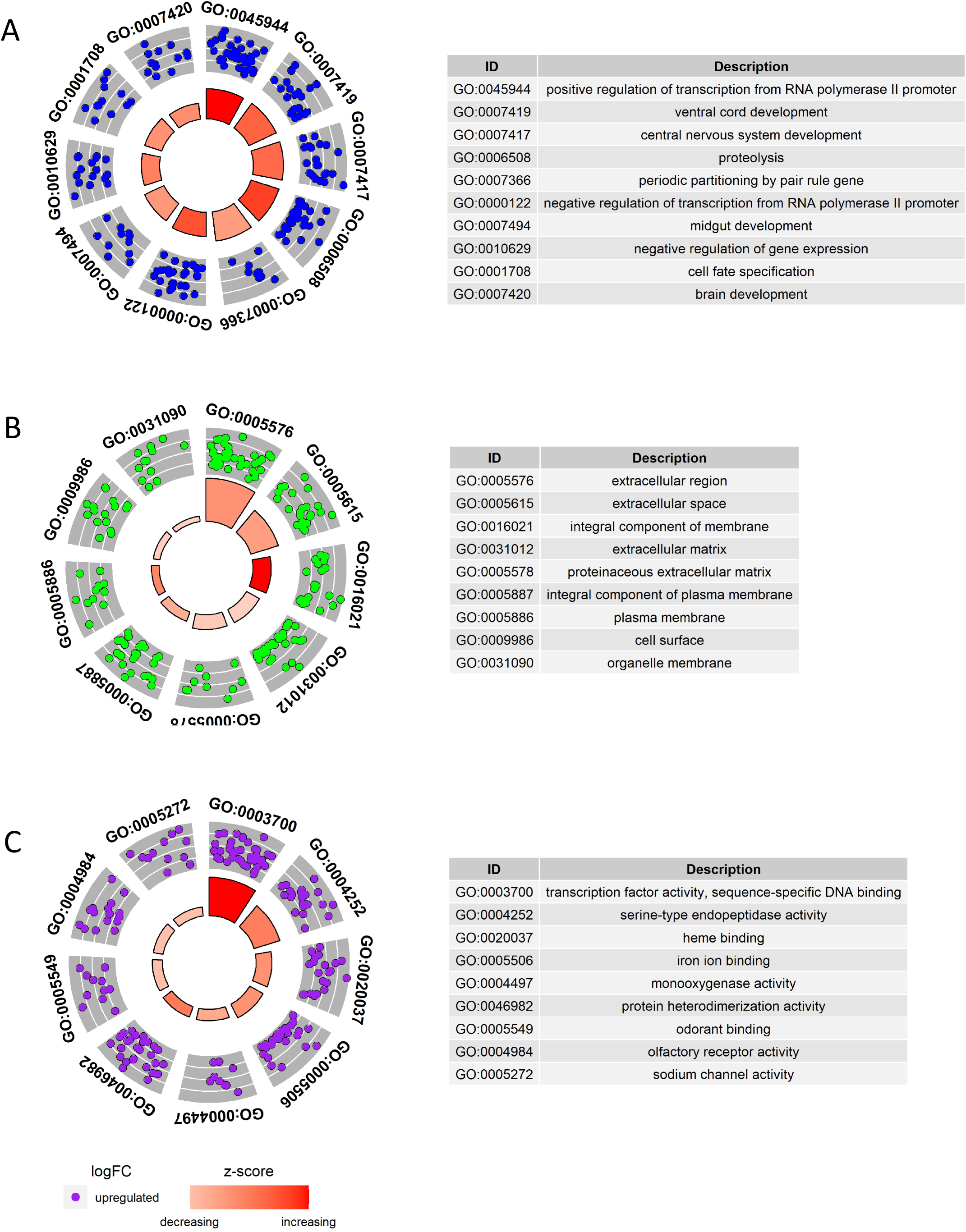
GO analysis of St5-only genes (Class 4). See the legend to Figure 3 for a description of the bars and circles on each panel. A) Biological Process GO terms. B) Cellular Component GO terms. C) Molecular Function GO terms.

#### Class 1: Stage 2-only

Our first group of genes have maternally supplied transcripts that are completely degraded by stage 5. They form the numerically smallest class (241 genes), and an analysis using DAVID did not show significant enrichment for any GO terms when compared to the full set of embryonically represented genes (Table S2). In other words, genes with transcripts that are completed degraded in early embryogenesis appear to be randomly distributed across functional categories and cellular components. We have previously shown that the St2-only state is rarely conserved across species (Atallah & Lott, 2018) for a given gene. It is possible that maternal deposition, where transcripts are dumped in the egg by the nurse cells, sometimes introduces developmental “noise” (the unnecessary deposition of RNA transcripts), and that the embryo compensates by degrading those transcripts.

#### Class 2: Stage 2 higher, stage 5 lower

These genes are represented at both stages, with lower levels at the later stage, and have maternally-supplied transcripts that are partially degraded by stage 5. (It is impossible to rule out that some of these genes may be zygotically transcribed, with the transcription compensating for at least some of the degradation.) We found 728 genes in this category with significant differential expression. We found the identified *D. melanogaster* orthologs to be enriched in the GO terms “mitochondrion”, “mitochondrial matrix” and “metabolic pathways”, although there was no significant enrichment for any specific pathway (Table S2). Furthermore, there was no significant enrichment of any GO terms in the Biological Process (BP), Cellular Component (CC) or Molecular Function (MF) categories. As with Class 1, genes with partially degraded transcripts are not limited to a specific subset of categories,

#### Class 3: Stage 2 lower, stage 5 higher

This class of genes include those that are represented at both stages, with levels that increase during development, and are believed to be both maternally supplied and zygotically transcribed. We found 1,364 genes in this category with significant differential expression between the two stages. They were enriched in products that play roles in protein and mRNA binding (Figure 3). Cellular components are localized to the plasma membrane and cytoplasm. They are also enriched in components of the Hippo and Notch signaling pathways, and have functions in RNA transport and ubiquitin-mediated proteolysis. Many of these genes encode protein kinases. Although genes in the Wnt signaling pathway were not significantly enriched, enrichment was seen for the InterPro (Hunter et al., 2009) term “armadillo-type fold”. Tandem armadillo repeats are found in many proteins involved in Wnt signaling. Signaling pathways play key roles during cellularization (stage 5), and segment polarity genes include components of Wnt signaling. The provision of these components through two mechanisms (maternal deposition and zygotic transcription) could be a form of redundancy to increase developmental robustness (Mestek Boukhibar & Barkoulas, 2016).

#### Class 4: Stage 5-only

We found 880 zygotic-only genes (represented only at St5). They encompass a range of important players in early development, and include genes involved in patterning, development of the central nervous system (including the brain) and several other ontogenetic processes (Figure 4A). Since St5 is the period when cellularization occurs, it is not surprising that more genes map to the GO term “integral component of membrane” than to any other cellular component (Figure 4B). They include genes encoding proteins localized to the plasma cell membrane and organelle membranes. Many of the gene products are also part of the extracellular matrix.

Not surprisingly, transcription factors, which activate patterning cascades and jump-start development, are by far the most strongly overrepresented molecular function class of St5-only genes, both in terms of the number of genes mapping to this category and the negative logarithm of the q-value (Figure 4C). Other zygotic-only genes have serine-type endopeptidase activity (GO term 0004252 in in Figure 4C); these include *masquerade*, which has a known role axonal development, and *modular serine protease* (modSP), which is involved in the immune system. St5-only genes also include heme-binding enzymes with oxidoreductive activity, including an array of cytochrome P450 enzymes (e.g. *phantom, disembodied, spookier* and many others).

### Some gain, and widespread loss, of gene expression in the *D. grimshawi* stage 5 embryo

The results of the differential expression analysis above were largely concordant with our previous findings for non-Hawaiian species (Atallah & Lott, 2018). We were curious, however, about any differences that might exist in gene representation in *D. grimshawi*, the first Hawaiian species with embryonic transcriptomes. When considering genes with *D. melanogaster* orthologs, we found more cases (101) of unique losses of stage 5 representation in *D. grimshawi* than in any other species (Figure 6).

**Figure 5.**
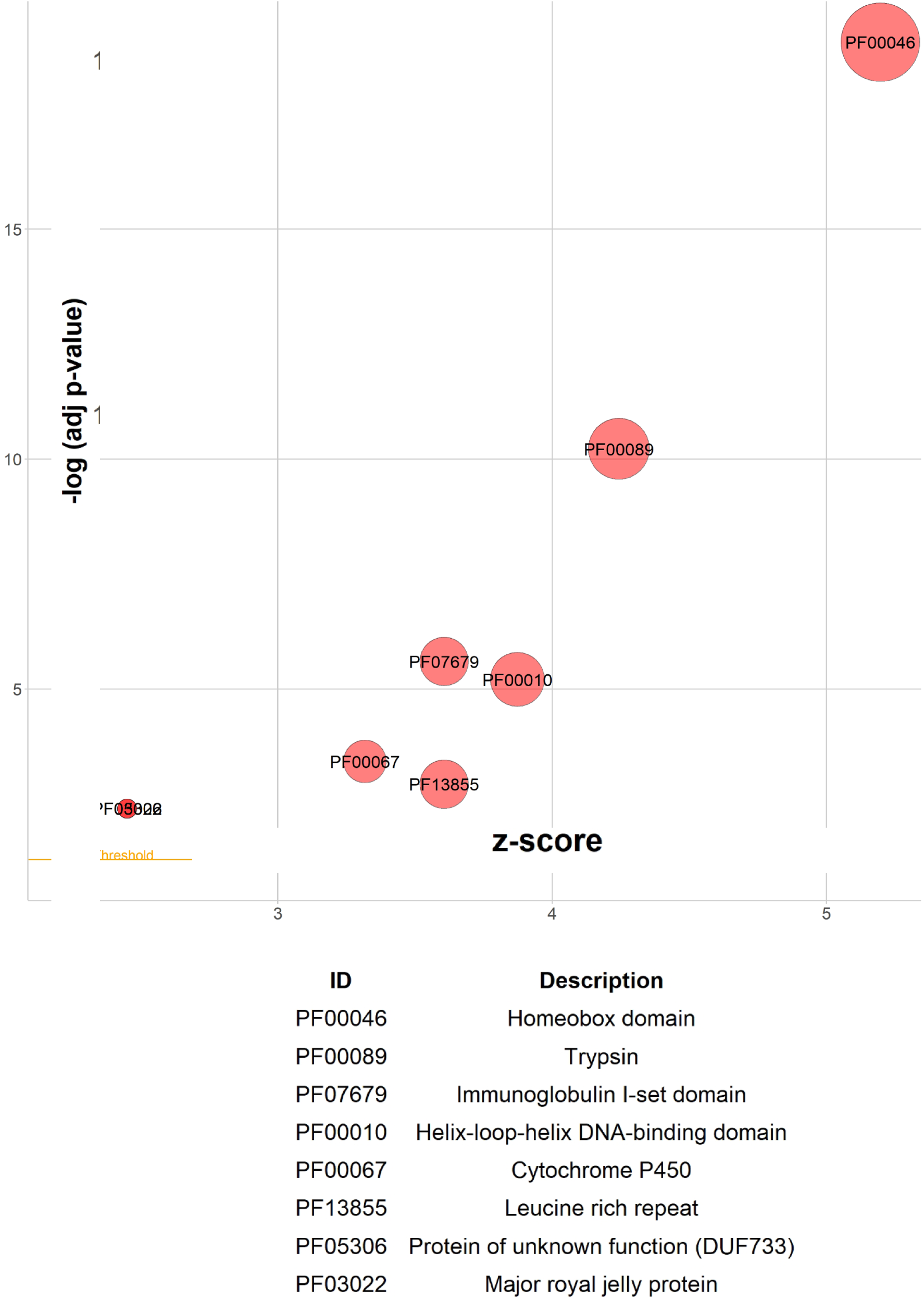
Protein Domain enrichment of Class 4 genes, using InterPro.

**Figure 6.**
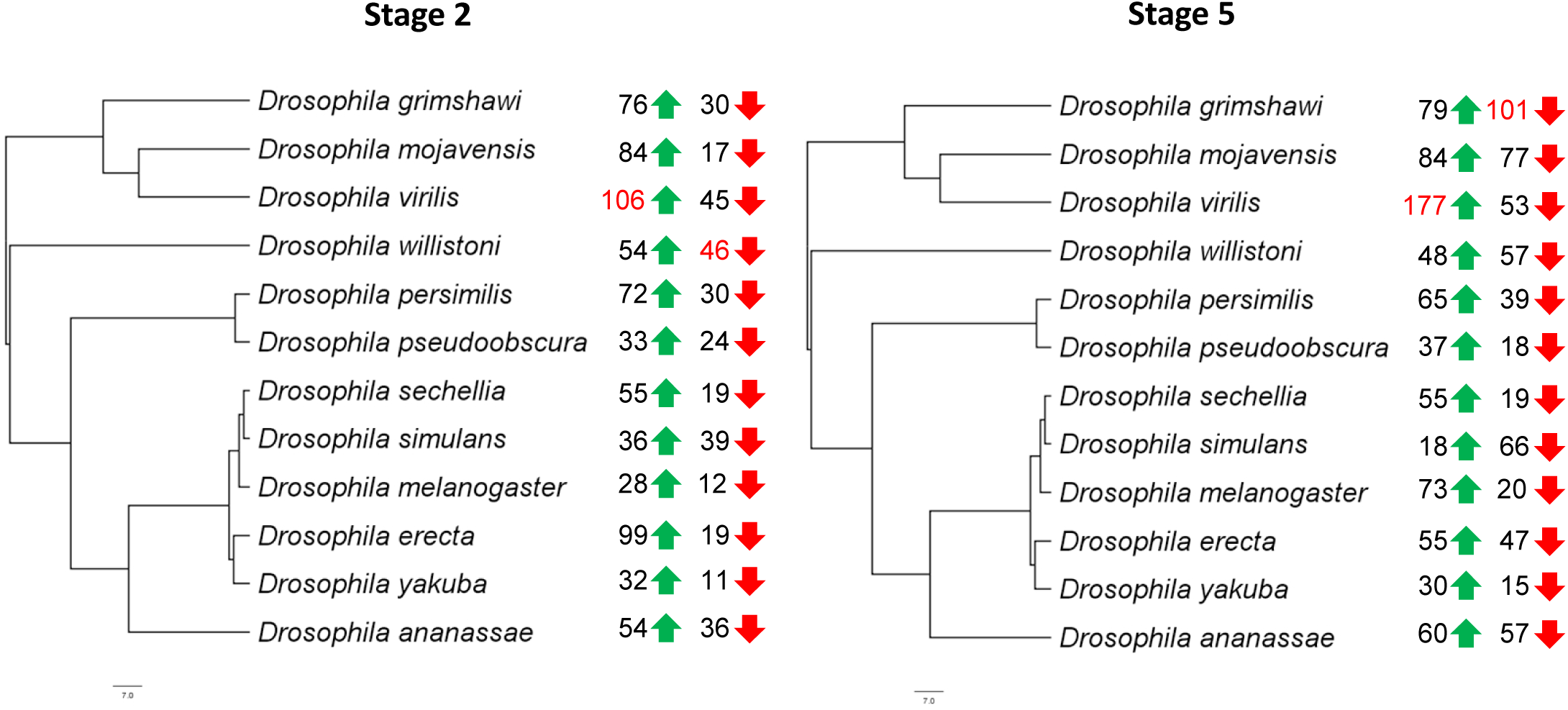
Numbers of unique gains (green arrows) and losses (red arrows) of St2 and St5 gene representation in each species. The analysis was conducted with 15 *Drosophila* species, but only the 12 originally sequenced species are shown (*D. mauritiana, D. santomea* and *D. miranda* are not shown). *D. virilis* shows the largest number of gains at both St2 and St5, while *D. willistoni* shows the largest number of losses at stage 2, and *D. grimshawi* shows the largest number of losses at stage 5.

Among the most interesting losses of St5 gene representation are the homeotic selector (Hox) genes, the downstream component of the anterior-posterior (AP) patterning pathway. As can be seen in Figure 7, multiple Hox genes are expressed by stage 5 in all species except *D. grimshawi*. We confirmed that components of the AP cascade upstream of the Hox genes, including the segment polarity, pair-rule and gap genes, are all expressed by stage 5 in *D. grimshawi* (Table S3).

**Figure 7.**
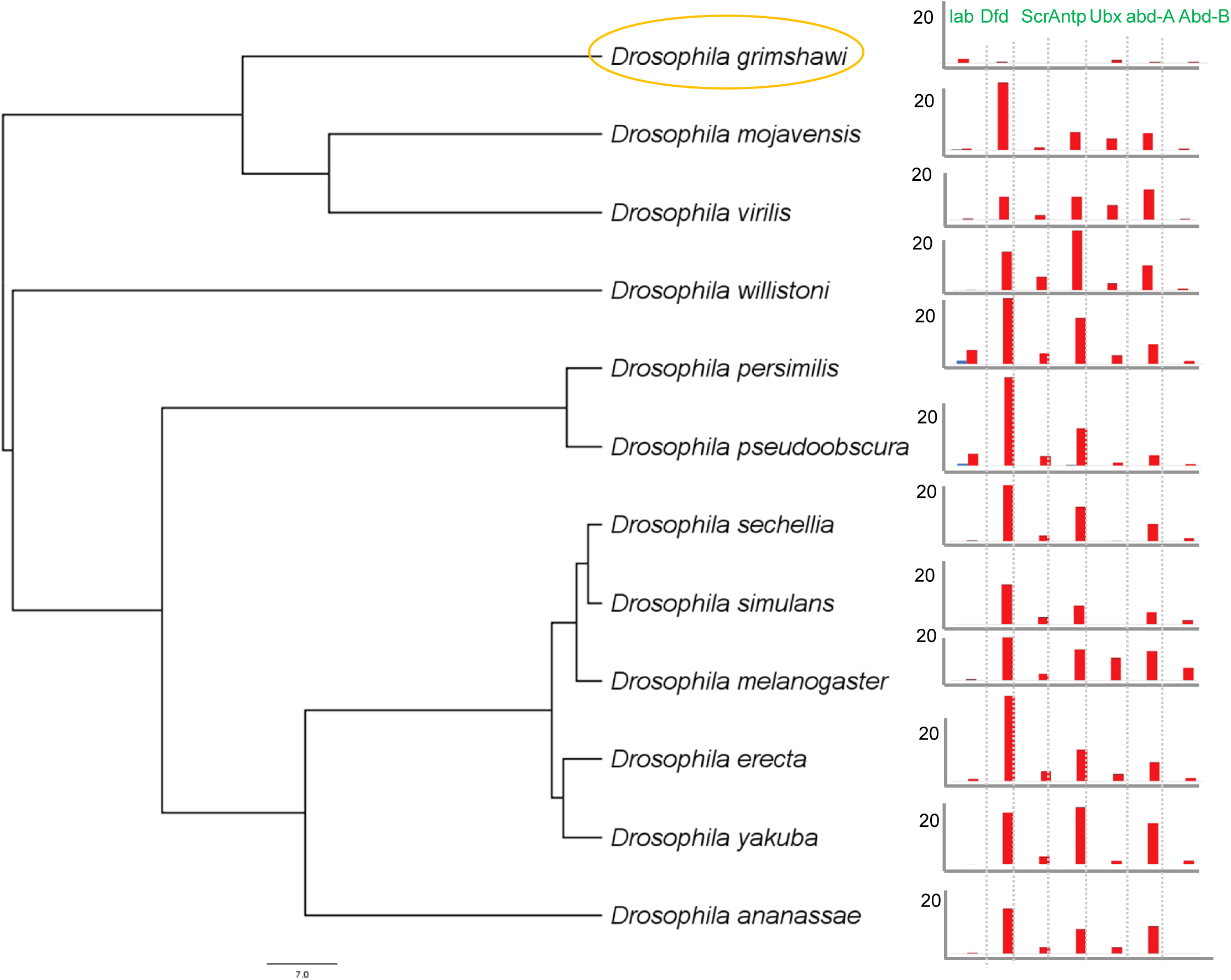
Loss of St5 representation of homeotic selector gene transcripts in *D. grimshawi*. St2 FPFM levels (close to 0 across all species) are shown in blue and St5 levels in red.

There are also a total of 79 genes with unique gains of representation (Figure 6). We took a closer look at the 35 genes with transcript abundance three times our threshold (FPKM > 3). Within this smaller set, 8 genes have named *D. melanogaster* orthologs, and are listed in Table S4. They include *Lim3*, a zinc finger transcription factor with a homeobox domain with a role in motor-neuron development. Another gene, *multiple wing hairs* (*mwh*), a downstream component of the planar cell polarity pathway, is represented in at both St2 and St5 in *D. grimshawi*, but not in any other species we have examined.

### Strong expression of unannotated genes in *D. grimshawi*

RNA-Seq bioinformatics methods that include the discovery of new isoforms, such as the Tuxedo suite, often uncover previously unannotated genes. In *D. grimshawi*, we find more unannotated transcripts with high stage 5 embryonic mRNA levels (FPKM > 3) than in any other species we have previously studied (Figure 7A). This finding could be partially due to the relatively poor annotation of the genome of this Hawaiian species (although the *D. grimshawi* genome annotation has improved markedly in recent years (Yang et al., 2018)). However, we were interested in knowing whether some of these transcripts could belong to putative taxonomically-restricted protein-coding genes.

A previous study in Drosophilidae (Heames et al., 2020) found unannotated genes in the family to be relatively short, coding for a median peptide length of only 81 amino acids. Using TransDecoder (Haas et al., 2013), we identified the longest complete open reading frame (ORF) (if any) in each of the 2,365 unannotated transcripts, and found 969 transcripts with an ORF of at least 50 amino acids. We used BLASTP (Altschul et al., 1990) to determine whether these putative peptides had homologs in the annotated genomes or embryonic transcriptomes of other *Drosophila* species. A total of 854 of the 969 ORFs had no identifiable orthologs. Of these, we chose to focus on the 301 transcripts with an FPKM above our threshold of 1 at Stage 2, Stage 5, or both.

### A subset of unannotated genes may have been generated de novo from non-coding sequence

Taxonomically restricted genes may have either originated from ancestral genes (e.g. through divergence beyond recognition from an ortholog, or through gene fission or fusion) or emerged de novo from non-coding sequence (Van Oss & Carvunis, 2019). Examples of de novo gene evolution, which at one time was dismissed as highly improbable, have fascinated researchers in recent years (Klasberg et al., 2018; Neme & Tautz, 2016; Zhao et al., 2014). Researchers frequently use TBLASTN to identify putative intergenic or intronic regions in other species that orphan genes might have arisen from (Lu et al., 2017; Sun et al., 2015). We adopted a similar approach, aided by the recent publication (B. Y. Kim et al., 2021) of new *Drosophila* genome assemblies (as yet unannotated), generated through Oxford Nanopore long-read sequencing, that included two additional Hawaiian picture-wing species (*Drosophila murphyi* and *Drosophila sproati*) along with 4 *Scaptomyza* species (*Scaptomyza graminum, Scaptomyza hsui, Scaptomya montana* and *Scaptomyza pallida*.)

Of the 114 transcripts with TBLASTN hits in one or more of the non-Hawaiian annotated genomes, the vast majority of the top hits (95, or 85%) are in intergenic or intronic regions in all species, suggesting that they may have arisen de novo from non-coding sequence (Figure 9A). Interestingly (Figure 8B), 97% of the 301 transcripts (291) had TBLASTN hits in either *D. murphyi* or *D. sproati* (the other two picture-wing Hawaiian *Drosophila* species). Only 47% had hits in the genomes of one of the four species in the *Scaptomyza* lineage, and 35% in the *Drosophila* subgenus (*D. virilis* or *D. mojavensis*).

**Figure 8.**
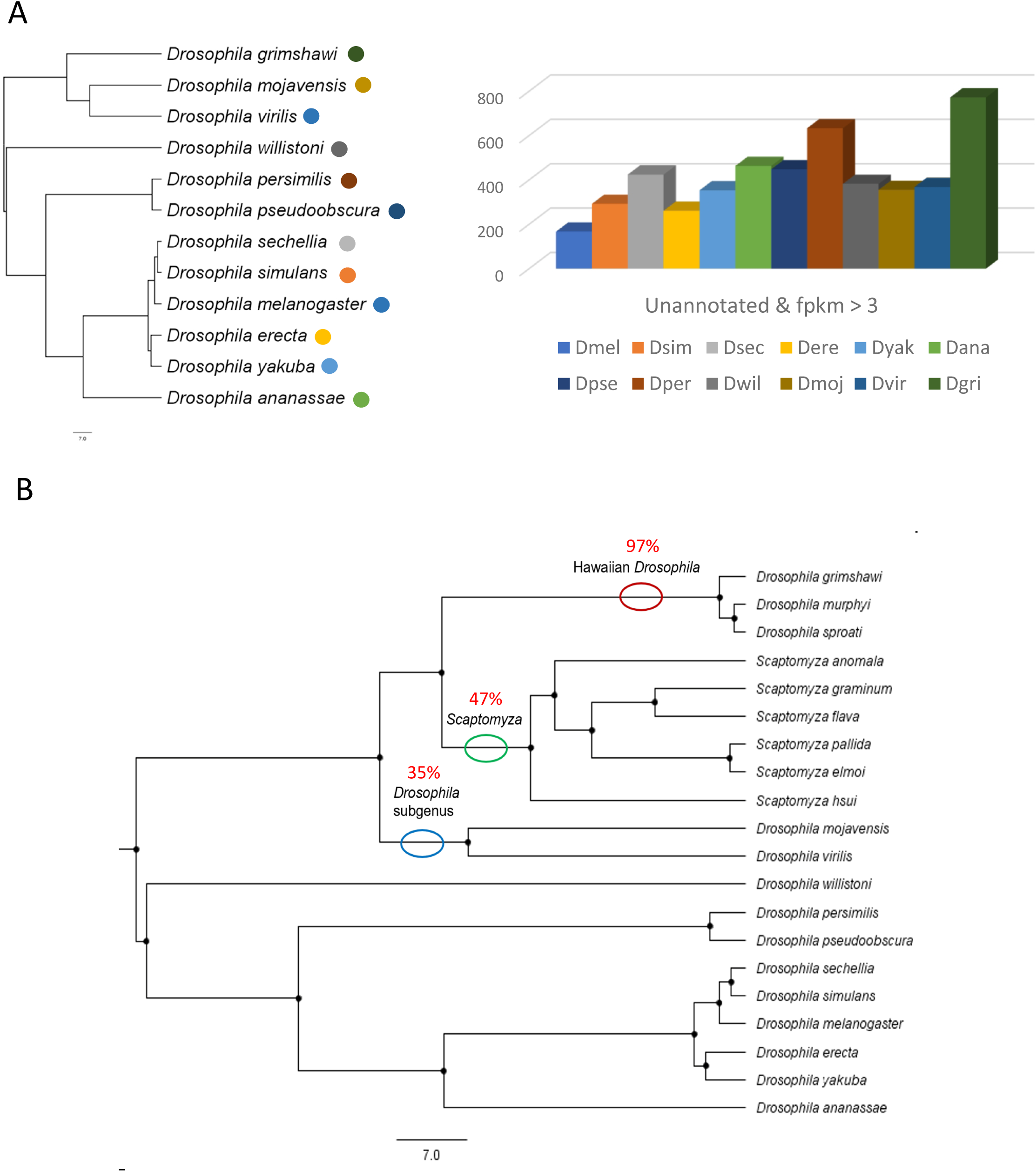
Unannotated genes in *D. grimshawi* show high expression levels, with a subset showing evidence of de novo evolution. A) More *D. grimshawi* unannotated genes are represented at high levels at stage 5 (FPKM > 3) than for any of the other species we have examined. B) Almost all identified ORFs in our candidate set of 301 *D. grimshawi* unannotated transcripts have TBLASTN hits in genomes of at least one other picture-wing Hawaiian *Drosophila* species, while slightly under half have hits in a *Scaptomyza* genome, and about a third have hits in *D. virilis* or *D. mojavensis*.

**Figure 9.**
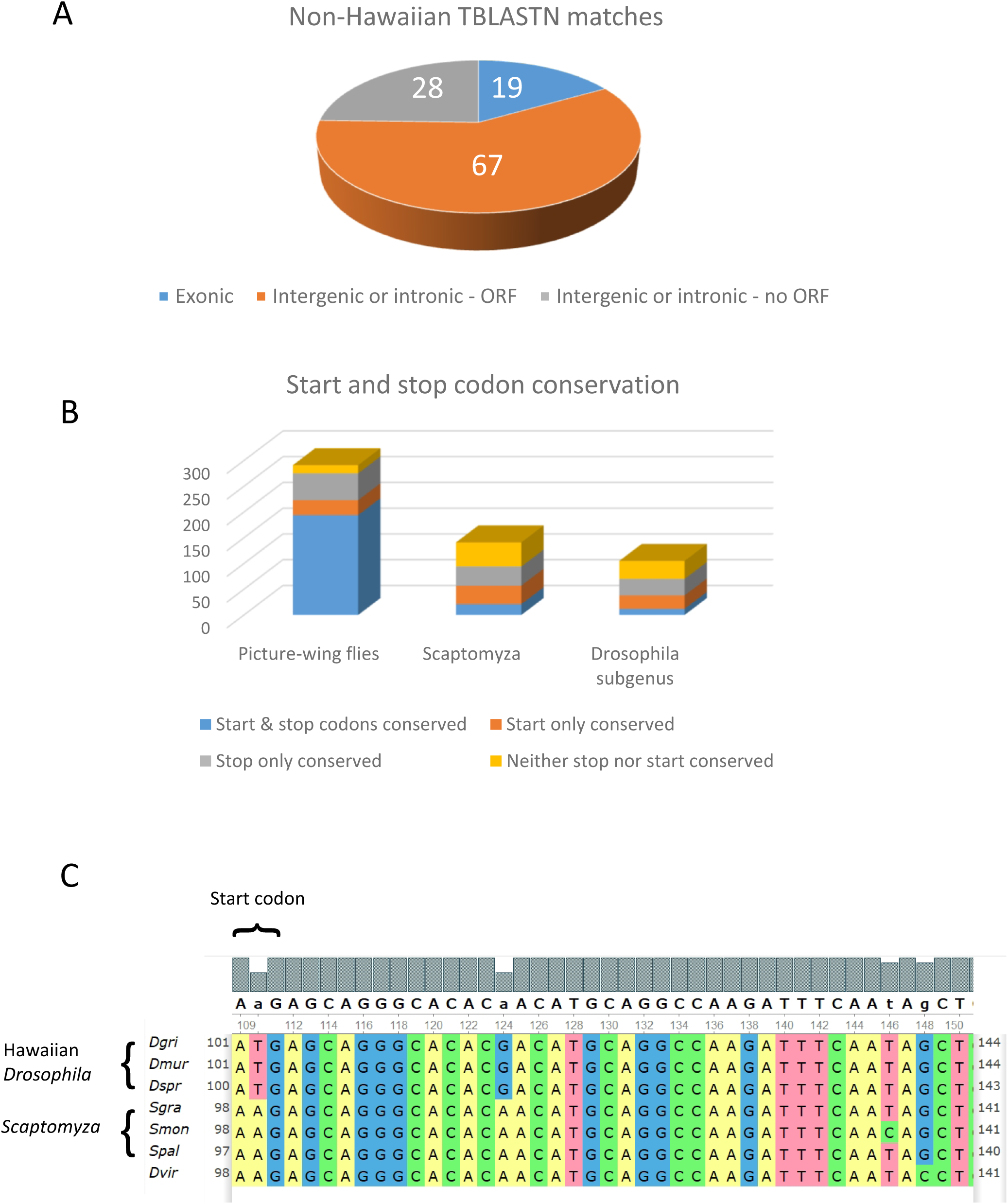
Putative de novo genes may have arisen from pre-existing ORFs or been generated from non-coding sequence. A) TBLASTN hits of possible *D. grimshawi* de novo transcripts (from the candidate set of 301) in non-Hawaiian species are overwhelmingly in intergenic or intronic regions, with a pre-existing ORF identified in the majority of cases. B) The start and stop codons of identified ORFs in *D. grimshawi* unannotated transcripts are highly conserved in other picture-wing species. Conservation is lower in *Scaptomyza* and the *Drosophila* subgenus. C) An example of a new ORF in a picture-wing Hawaiian *Drosophila* species. In the unannotated *D. grimshawi* transcript TCONS_41457, a start codon evolved from a single nucleotide change in the ancestor of the picture-wing Hawaiian *Drosophila* species. The alignment was generated using CLUSTALW.

Of the transcripts with noncoding TBLASTN hits in annotated non-Hawaiian species genomes, 67 out of 95 (71%) matched regions that contain an ORF in at least one of the fly genomes (Figure 9A), possibly lending support to an ORF-first model of de novo gene evolution. Alignments with CLUSTALW showed that in most cases both the start and stop codons were conserved in at least one of the other picture-wing *Drosophila* species (Figure 9B). The same was true far less frequently in *Scaptomyza* and the *Drosophila* subgenus species. We also found cases, however, where a start or stop codon may have been generated from non-coding sequence through a single-nucleotide substitution (Figure 9C). Our results suggest that a subset of unannotated genes with embryonic mRNA representation may have been generated de novo from non-coding sequence.

## DISCUSSION

Island endemics are excellent systems for exploring the evolution of novelty. Most famously, the flora and fauna of the Galapagos islands were critical to Charles Darwin’s (Darwin, 1845) development of the theory of natural selection. More recently, the evolution of Anolis lizards on the Greater and Lesser Antilles has been analyzed rigorously (Corbett-Detig et al., 2020; Mahler et al., 2010, 2016). Islands such as the Hawaiian archipelago, which arose within the past five million years through volcanic eruptions, presented pristine, untouched environments for early colonists with untapped ecological opportunities (Whittaker et al., 2017). Evolution on these types of islands proceeds far more rapidly than on the mainland (Losos & Ricklefs, 2009), leading to diversity which often exceeds those of continental species.

As close relatives of the model organism *D. melanogaster*, the Hawaiian *Drosophila* are a particularly valuable model for studying the evolutionary genetics of island radiations. Research on this clade has slowed, however, in recent decades (O’Grady & DeSalle, 2018). We present here the first transcriptomic comparison in a Hawaiian fly of two iconic stages in embryogenesis that we know to be highly conserved across Drosophilidae (Kuntz & Eisen, 2014). We find that while *D. grimshawi*’s early embryonic transcriptome at these stages is similar in many ways to other non-Hawaiian species, there are distinct differences, with numerous losses of orthologous gene representation. Most notably, while the Hox genes are represented by stage 5 in every other species we have examined, we find no evidence of this in *D. grimshawi*. Future studies will be necessary to determine when homeotic selector genes are activated in this species, and whether the apparent delay in Hox activation is also seen in other Hawaiian flies.

Genes with zygotic (i.e. stage 5) expression in *D. grimshawi*, but not in other species (Table S4), are excellent candidates for functional knock-out experiments. We are currently developing protocols for targeted mutagenesis in *D. grimshawi* using the CRISPR-Cas9 system (Doudna & Charpentier, 2014; Jinek et al., 2012). The distinct advantage of examining the early embryo is that the effects of mutations can be assessed at the beginning of development. While in the long-term we would like to examine the activity and functions of genes active in larval and pupal discs of structures such as wings and legs, which have diverged markedly in Hawaiian species (Edwards et al., 2007; Stark & O’Grady, 2010), these goals will be more challenging to achieve, because the role of a gene early in development may mask its function at a later stage (e.g. if the gene is embryonically lethal, it is more difficult to determine its later role in imaginal discs). Such research may require precise editing of specific enhancers.

Gains and losses of expression of genes with orthologs in other species, including the model fly *D. melanogaster*, could be examples of co-option of a conserved genetic toolkit (Carroll, 2008), although the functional experiments described above will be necessary for investigating this possibility. While the re-use of existing genes is thought to play an important role in developmental evolution (Stern, 2011), attention has turned more recently to the role of taxonomically restricted genes (Johnson, 2018; McLysaght & Guerzoni, 2015; Van Oss & Carvunis, 2019). Much of the work investigating de novo gene evolution has focused on transcripts expressed in the testis (Lange et al., 2021; Witt et al., 2019; Zhao et al., 2014), where evolutionary turnover occurs rapidly (Jagadeeshan & Singh, 2005; Mohammed et al., 2014), with fewer studies looking at genes expressed in relatively slowly-evolving systems such as embryogenesis. In this study, we were interested in the fact that more unannotated genes were expressed in *D. grimshawi* at St5 than in any of the 14 other species we had previously examined. While we had speculated (Atallah & Lott, 2018) that unannotated genes might be taxonomically restricted, we had not examined this possibility further, and it is possible that many unannotated genes are simply a result of poor genome annotation. In this study, we identified a total of 95 embryonic *D. grimshawi* transcripts with open reading frames that had homologous sequence matches in intergenic or intronic regions in a non-Hawaiian species (Figure 9A), suggesting that they may have originated de novo, either in the Hawaiian *Drosophila* lineage or the ancestor of Hawaiian *Drosophila* and *Scaptomyza*.

Two models exist to explain the emergence of de novo protein-coding genes (McLysaght & Guerzoni, 2015). In the “ORF-first” model, a pre-existing ORF is expressed through regulatory element evolution. In a second model, a non-coding RNA acquires an ORF (Xie et al., 2012). Analyzing embryonic transcriptomes from other Hawaiian *Drosophila* and *Scaptomyza* species, where we often found ORFs in regions that were homologous to the unannotated expressed sequences in *D. grimshawi*, will be necessary to distinguish between these models. As with co-opted genes, targeted mutagenesis of putative de novo genes will be important for determining whether they have acquired novel functions in the early embryo.

## ACKNOWLEDGEMENTS

We would like to thank Delmy Urbina, Jacob Michilak and Bianca Canizares for assistance with the experiments. Funding for this work was provided to Joel Atallah from the Louisiana Board of Regents (grant number LEQSF(2017-20)-RD-A-26).

## DATA AVAILABILITY STATEMENT

## SUPPORTING INFORMATION – TABLE LEGENDS

**Table S1**. Gene FPKM levels in previously analyzed species (Atallah & Lott, 2018) and *D. grimshawi*. Dmel = *Drosophila melanogaster*, Dsim = *Drosophila simulans*, Dsec = *Drosophila sechellia*, Dmau = *D. mauritiana*, Dyak = *Drosophila yakuba*, Dsan = *Drosophila santomea*, Dere = *Drosophila erecta*, Dana = *Drosophila anananasse*, Dpse = *Drosophila pseudoobscura*, Dper = *Drosophila persimilis*, Dmir = *Drosophila miranda*, Dwil = *Drosophila willistoni*, Dmoj = *Drosophila mojavensis*, Dvir = *Drosophila virilis*, Dgri = *Drosophila grimshawi*.

**Table S2**. Gene Ontology analysis using DAVID (Huang et al., 2009a, 2009a).

**Table S3**. Segmentation gene FPKM levels at St2 and St5. *D. grimshawi* and 14 other (Atallah & Lott, 2018) species are shown.

**Table S4**. List of genes with named *D. melanogaster* one-to-one orthologs that are strongly represented at St5 in *D. grimshawi* (FPKM > 3) and not represented at that stage in 14 other *Drosophila* species.

